# Comparative Analysis of Gene Expression Analysis Methods for RNA In Situ Hybridization Images

**DOI:** 10.1101/2024.02.25.581931

**Authors:** Valeria Ariotta, Eros Azzalini, Vincenzo Canzonieri, Sampsa Hautaniemi, Serena Bonin

**Affiliations:** Research Program in Systems Oncology, Research Programs Unit, Faculty of Medicine, University of Helsinki, 00014 Helsinki, Finland; Department of Medical Sciences (DSM), University of Trieste, Trieste, 34149, Italy; Pathology Unit, Centro di Riferimento Oncologico (CRO) IRCCS, Aviano-National Cancer Institute, Pordenone, 33081, Italy

**Keywords:** RNA, ISH, RT-ddPCR, automated quantification, FFPE

## Abstract

Gene expression analysis is pivotal in cancer research and clinical practice. While traditional methods lack spatial context, RNA in situ hybridization (RNA-ISH) is a powerful technique that retains spatial tissue information. Here, we investigated RNAscope score, RT-droplet digital PCR (RT-ddPCR), and automated QuantISH and QuPath in quantifying RNA-ISH expression values from formalin-fixed paraffin-embedded samples. We compared the methods using high-grade serous ovarian carcinoma samples, focusing on *CCNE1*, *WFDC2*, and *PPIB* genes. Our findings demonstrate good concordance between automated methods and RNAscope, with RT-ddPCR showing less concordance. We conclude that QuantISH exhibits robust performance, even for low-expressed genes like *CCNE1*, showcasing its modular design and enhancing accessibility as a viable alternative for gene expression analysis.

## Introduction

Reliably estimating gene expression values is of paramount importance in cancer research and various clinical tasks, including diagnostic and prognostic (Golub et al., 1999; Kamel & Al-Amodi, 2017; Narrandes & Xu, 2018; van ‘t Veer et al., 2002). Over the past decades, the abundance of mRNA molecules has been detected using RT-PCR, microarrays, and sequencing (Garber et al., 2011; Macgregor & Squire, 2002; Taylor et al., 2017). While these technologies resulted in several advances (Finotello & Di Camillo, 2015; Fryer et al., 2002; Jozefczuk & Adjaye, 2011; Rai et al., 2018), they measure gene expression from bulk or single cells without considering the spatial context. In contrast, in situ hybridization (ISH) technologies, such as the RNAscope assay (Atout et al., 2022; Hua et al., 2018; Olmedillas-López et al., 2017; Wang et al., 2012) and spatial transcriptomics (Rao et al., 2021; Tian et al., 2023), are capable of quantifying gene expression in single cells in their spatial context (Burgess, 2019; Chen et al., 2015; Jamalzadeh et al., 2022). The spatial preserving transcriptomics technologies not only offer an enhanced alternative for conventional biomarker detection but also represent a substantial transformation in research disciplines such as oncology and neuroscience (Baleriola et al., 2014; Bingham et al., 2017; Kunju et al., 2014; Wang et al., 2012).

In the analysis of RNA in situ hybridization (RNA-ISH) data, diverse approaches exist, including semi-quantitative assessment on the standard microscope, where human involvement remains essential for result interpretation, to automated image analysis that allows the streamlining of the analytic process, offering a standardized and efficient pathway to quantify and characterize RNA expression patterns. In particular, digital image analysis can be performed through dedicated software such as QuPath (Bankhead et al., 2017) or fully automated pipelines such as QuantISH (Jamalzadeh et al., 2022), SMART-Q (Yang et al., 2020), and dotdotdot (Maynard et al., 2020).

Herein, we studied the expression levels of oncogene *CCNE1*, diagnosis biomarker *WFDC2,* and one housekeeping gene (*PPIB*) on formalin-fixed paraffin-embedded (FFPE) samples from patients with ovarian high-grade serous carcinoma (HGSC). HGSC is the most prevalent subtype of tubo-ovarian cancer and is known for its low five-year survival rate of only 40% (Fabbro et al., 2020). *CCNE1* amplification, occurring in 15-20% of HGSC cases (Bell et al., 2011; Jamalzadeh et al., 2022; Marone et al., 1998), has been linked to HGSC patient survival (Chan et al., 2020; Jamalzadeh et al., 2022; Stronach et al., 2018). *WFDC2* is a well-established ovarian cancer biomarker (AlSomairi et al., 2024; Braicu et al., 2022; Utkarsh et al., 2023) and is also associated with HGSC patient survival (James et al., 2022). *PPIB* was selected as a control gene, and it is already used in several ovarian cancer studies by RNAscope assay and RT-PCR analysis (Desbois et al., 2020; Pan & Ma, 2020). The expressions of these genes were assessed using RT-droplet digital PCR (ddPCR) and on matched tissue microarrays (TMA) slides by RNAscope assay, analyzing RNA signal both digitally (QuantISH and QuPath pipelines) and in a semi-quantitative fashion (RNAscope score).

We conduct a comparative analysis of the four methods utilized for gene expression assessment to investigate the reliability, concordance, and limitations of the approaches. The justification for selecting these four methods is as follows: i) RNAscope and RT-ddPCR represent widely recognized probe-based techniques commonly employed in gene expression analysis; ii) QuantISH and the QuPath pipeline constitute, to our knowledge, the only two available open-source pipelines for RNA-ISH analysis.

## Results

The average expression levels of individual genes were evaluated using different methodologies: in situ visual assessment by RNAscope score, in situ digital assessment by QuantISH and QuPath pipelines, and RT-ddPCR analysis on bulk tissue. The results for each method are summarized in Table 1. Upon comparison, all methods produce a similar order of gene expression, with *WFDC2* having the highest expression, *CCNE1* the lowest, and *PPIB* in between (Friedman test; p < 0.0001). Gene pairwise comparisons returned significant p-values in all methodologies (Wilcoxon paired test; p < 0.0001). These findings emphasize the robustness of computational methods in effectively distinguishing varying expression levels across genes, regardless of their differing magnitudes of expression.

**Table 1.** Descriptive statistics of gene expression values for each method.

|  |  | CCNE1 | PPIB | WFDC2 |
| --- | --- | --- | --- | --- |
| log10 QuantISH signal | mean | -2,9 | -2,1 | -1,7 |
|  | min | -5,1 | -3,8 | -4,6 |
|  | max | -1,3 | -1,1 | -0,78 |
|  | CV | 24% | 27% | 38% |
| n° estimated spots<br>QuPath | mean | 3 | 12 | 22 |
|  | min | 0 | 1 | 0 |
|  | max | 21 | 35 | 59 |
|  | CV | 132% | 60% | 66% |
| RNAscope score (0-4) | mean | 1 | 2 | 3 |
|  | min | 0 | 0 | 0 |
|  | max | 4 | 4 | 4 |
|  | CV | 59% | 32% | 31% |
| log2 copies/μl RT-ddPCR | mean | 2,7 | 7,2 | 8,2 |
|  | min | -3,5 | 2 | 2,1 |
|  | max | 7,5 | 10 | 13 |
|  | CV | 78% | 25% | 28% |

### Accuracy of gene expression assessment in comparison to visual scoring

We next assessed the concordance between the RNAscope visual scoring system and the respective expression values quantified by QuantISH, QuPath, and RT-ddPCR for *CCNE1*, *PPIB*, and *WFDC2* genes, as illustrated in Figure 1. The RNAscope scoring system, known for its histological assessment, served as a benchmark for our comparative analysis, providing a standardized basis for evaluating the performance of different methodologies in distinguishing between these distinct scoring categories. The frequencies of RNAscope scores for each gene are represented in Supplementary Fig. 2.

**Figure 1.**
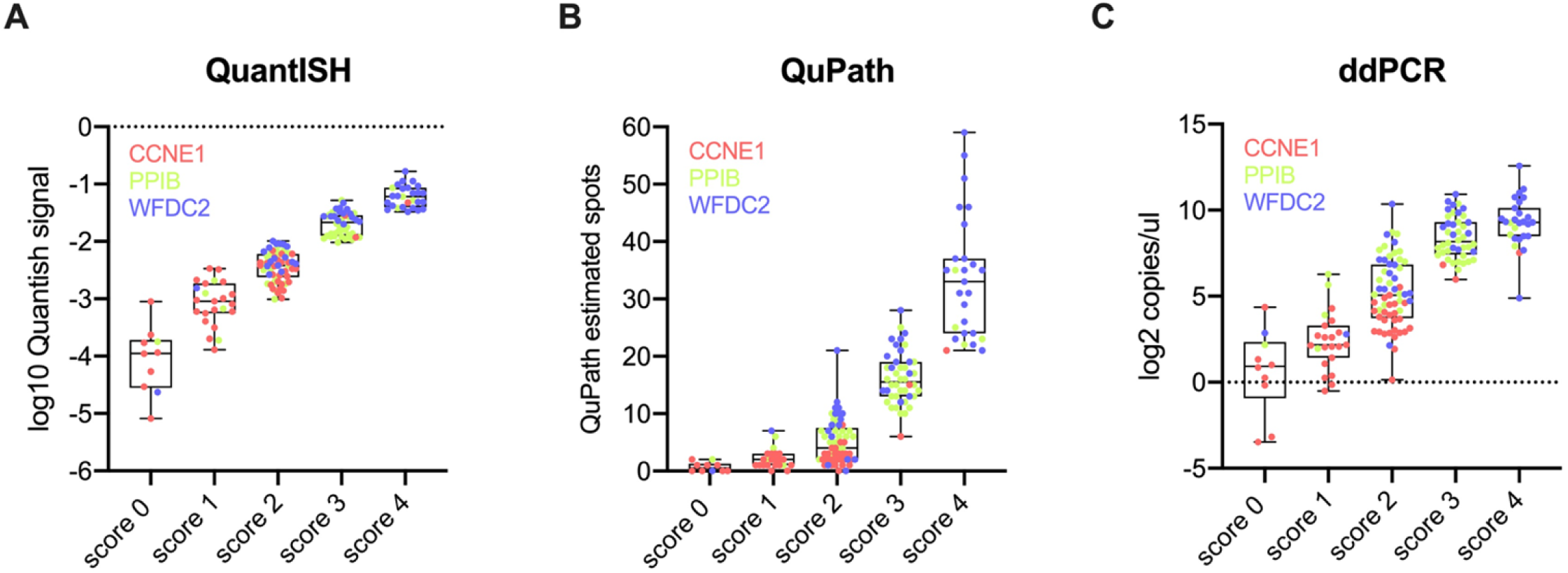
Boxplot representing the concordance between visual estimation by RNAscope score and gene expression level quantified by QuantISH (A), QuPath (B), and RT-ddPCR (C) for *CCNE1* (red dots), *PPIB* (green dots), and *WFDC2* (blue dots).

Overall, all methods were able to distinguish between any two scoring bins with high statistical significance (Mann-Whitney test, p < 0.0001). However, the mean difference between scores 0 and 1 was less pronounced for QuPath (p = 0.01) and RT-ddPCR (p = 0.02) than QuantISH. Additionally, for RT-ddPCR, the mean difference between scores 3 and 4 was less pronounced (p = 0.005).

### Inter-methods correlations

There was a high and statistically significant correlation among all methodologies (Fig. 2A). In particular, QuantISH and QuPath demonstrated a noteworthy alignment across all methodologies (mean ρ = 0.91 for QuantISH and 0.90 for QuPath), exhibiting a strong correlation with RNAscope visual assessment (ρ = 0.94 for QuantISH and 0.90 for QuPath). Conversely, RT-ddPCR showed slightly lower but still notably high correlation coefficients among the different methods (mean ρ = 0.85).

**Figure 2.**
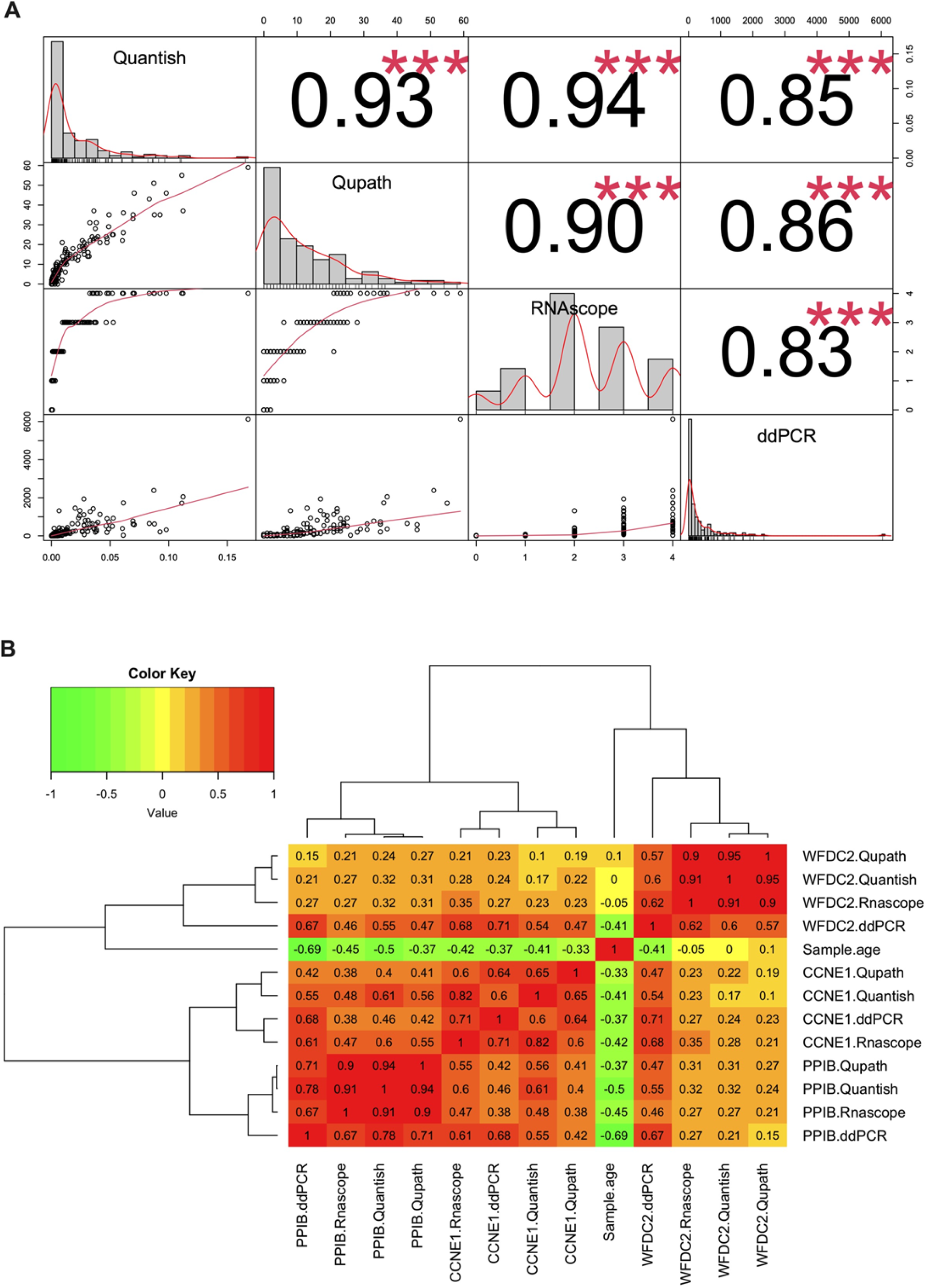
A) Scatter plot matrix for gene expression values obtained by RNAscope visual assessment, QuantISH, QuPath, and RT-ddPCR analysis of the 55 FFPE samples. In this analysis, the pairwise Spearman Rho coefficient was calculated. B) Heatmap with hierarchical clustering analysis for the Spearman correlation of transcript abundance as determined by RNAscope visual assessment, QuantISH, QuPath, and RT-ddPCR for all pair-wise combinations of *CCNE1*, *PPIB*, *WFDC2*, and sample age.

Upon examining individual genes more closely, we observed varying correlations among methods (Fig. 2B). The mean inter-method correlations for *CCNE1*, *PPIB*, and *WFDC2* were ρ = 0.67, 0.82, and 0.76, respectively. The *CCNE1* gene, which has low expression levels, showed the lowest correlation values compared to *PPIB* and *WFDC2*, which have moderate and high expression levels, respectively. QuantISH maintained high correlation values with the RNAscope score for all genes analyzed (mean ρ = 0.88, range 0.82-0.91), while QuPath and RT-ddPCR exhibited gene-specific variations (mean ρ = 0.80, range 0.60-0.90 and mean ρ = 0.67, range 0.62-0.71, respectively). QuPath exhibited lower concordance for the *CCNE1* gene, while RT-ddPCR exhibited lower concordance for *WFDC2*. Notably, RT-ddPCR consistently yielded lower correlation values than the other methods.

Observing the association between methods and block’s age - the time interval between surgery, when tissue block was made, and RNAscope assay date - we noted significantly higher correlation values for *PPIB*, a housekeeping gene, in comparison to *CCNE1* and *WFDC2* across all methods (Fig. 2B). Particularly striking was the pronounced drop-off in the housekeeping gene *PPIB* expression levels observed in RT-ddPCR (r = −0.69) compared to the in-situ hybridization methods (mean ρ = −0.44, range −0.37- −0.50), suggesting that the gene expression values based on RT-ddPCR are more sensitive to sample age due to nucleic acid degradation.

### Inter-methods correlations according to gene expression level

The variation in the level of agreement between QuPath, QuantISH, and RT-ddPCR was then analyzed within each RNAscope score as determined by visual assessment (Fig. 3). QuPath and QuantISH showed a high degree of agreement in the detection of spots without significant signal (score 0; ρ = 0.84), while an increase in correlation values was observed when moving from low to high expression levels, namely from score 1 (ρ = 0.44) to 4 (ρ = 0.71). On the contrary, RT-ddPCR showed higher concordance with both QuantISH and QuPath for scores 2-3 (medium level of expression), while for scores 0-1 (low expression) and score 4 (high expression), the correlation values were low.

**Figure 3.**
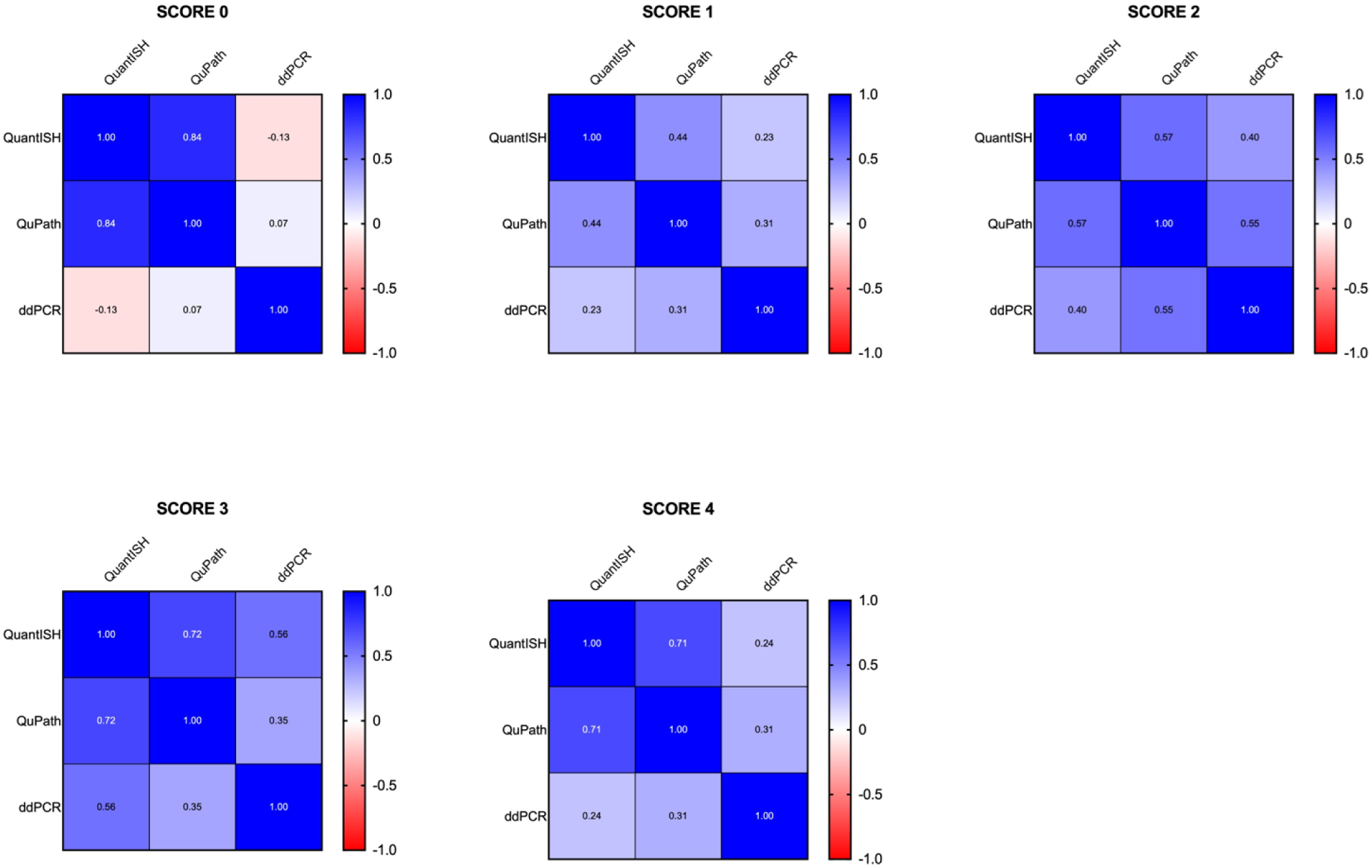
Correlation matrix for gene expression values obtained by QuantISH, QuPath, and RT-ddPCR according to the different RNAscope scores (0-4).

### Concordance between RNAscope scoring detected by visual estimation and Qupath analysis

To evaluate the concordance between the RNAscope visual scoring system and QuPath directly, we created RNAscope scoring groups based on the estimated number of spots in QuPath. For all genes, the Bland-Altman analysis resulted in a bias of −0.036 with a 95% limit of agreement ranging from −1.37 to 1.30 (Figure 4A). The pairwise comparison of visual and digital scoring showed 41% discordant cases (Fig. 4B); the highest discordance rates were detected for score 1 vs 2 (40%), followed by scores 3 vs 4 (33%), 2 vs 3 (12%), and 0 vs 1 (10%). Notably, the discordant cases were mainly represented in QuPath software by the estimated spots between 0 and 6 and, to a lesser extent, by the estimated spots greater than 15, as shown in Figure 4C. No significant difference in the proportion of concordant and discordant samples was observed when looking at individual genes (Chi-square test; p = 0.1) (Figure 4D). However, *CCNE1* had the highest discordance rate (51%), followed by *WFDC2* (40%) and *PPIB* (31%).

**Figure 4.**
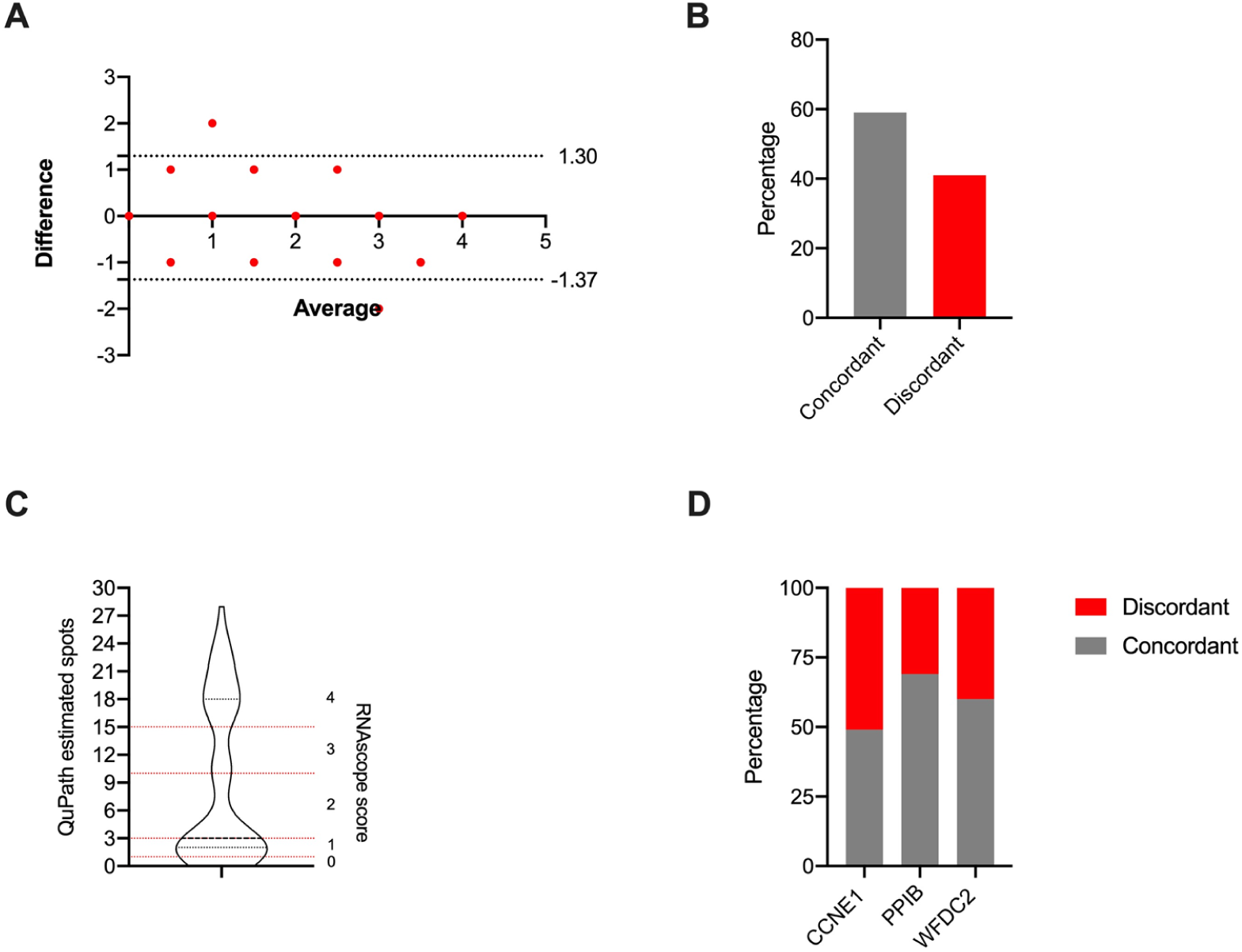
A) Bland-Altman plot representing the concordance between RNAscope scores detected by visual assessment and those derived from QuPath digital analysis. B) Percentage of cases sharing the same RNAscope scores between visual assessment and QuPath digital assessment. C) Violin plot representing the distribution of the estimated number of spots as detected by QuPath in discordant cases. D) Percentage of concordant and discordant cases by gene.

### Inter-methods correlations – Normalized data

The expression levels of *CCNE1* and *WFDC2* were normalized against those of *PPIB* to account for sample degradation. Subsequently, normalized data were used to conduct a Spearman correlation analysis to re-test inter-method agreement. Overall, all methods displayed a good level of correlation (Fig. 5A). QuantISH and QuPath exhibited the highest inter-method agreement (mean ρ = 0.85 and 0.84, respectively), followed by RT-ddPCR and RNAscope score (mean ρ = 0.81 and 0.78, respectively). Both in situ automated methods and RT-ddPCR showed a good to medium level of agreement with the RNAscope score (ρ = 0.84 for QuantISH, 0.76 for QuPath, and ρ = 0.74 for RT-ddPCR).

**Figure 5.**
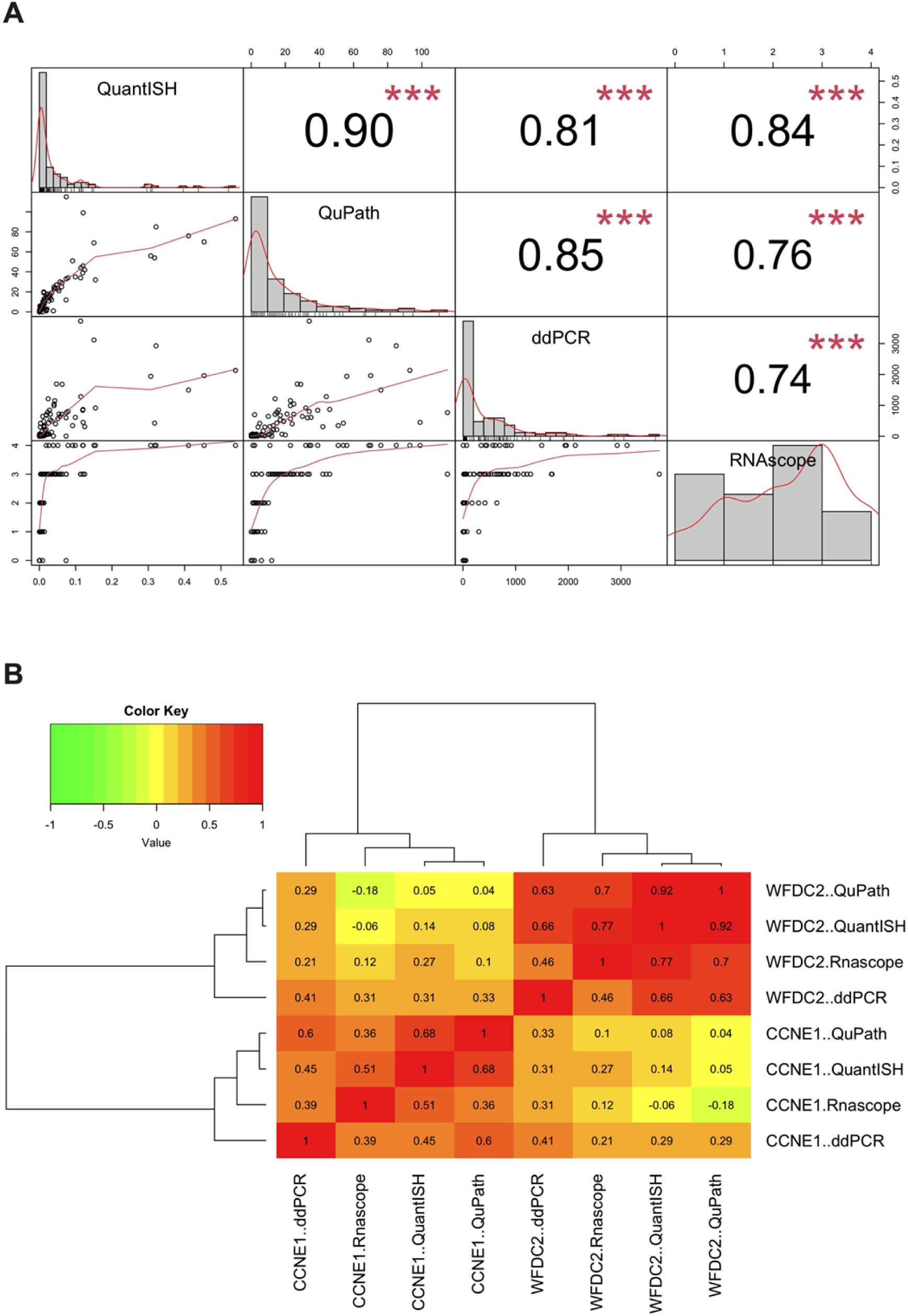
A) Scatter plot matrix for normalized gene expression values obtained by RNAscope visual assessment, QuantISH, QuPath, and RT-ddPCR analysis of the 55 FFPE samples. In this analysis, the pairwise Spearman Rho coefficient was calculated. B) Heatmap with hierarchical clustering analysis for the Spearman correlation of normalized transcript abundance as determined by RNAscope visual assessment, QuantISH, QuPath, and RT-ddPCR for all pair-wise combinations of *CCNE1* and *WFDC2* genes.

Both *CCNE1* and *WFDC2* displayed a medium level of concordance between methods (Fig.5B; mean ρ = 0.50 (*CCNE1*) and 0.69 (*WFDC2*)). However, QuantISH showed a higher correlation with the RNAscope score for the low-expressed gene *CCNE1* (ρ = 0.51) compared to QuPath and RT-ddPCR (ρ = 0.36 and 0.39 respectively). At high expression levels of *WFDC2*, both in situ digital methods performed significantly better than RT-ddPCR (ρ = 0.77 for QuantISH and 0.70 for QuPath, compared to 0.46 for RT-ddPCR).

## Discussion

We conducted a comparative analysis of four methods for gene expression assessment. Three methods involved quantifying RNA expression in TMA slides using visual assessment (RNAscope score) and automated image analysis (QuantISH and QuPath pipelines). The fourth method involved absolute target quantification using RT-ddPCR on matched bulk tissue. Specifically, our study aimed to highlight the practical utility of fully automated digital image analysis, demonstrating its potential as a viable alternative to the traditional RNAscope score approach and evaluating its concordance with a gold standard such as RT-ddPCR.

All methods used in this study were suitable for analyzing *CCNE1*, *PPIB*, and *WFDC2* gene expression. QuantISH and QuPath, showed a good and remarkably high correlation with the RNAscope visual score. However, some differences were observed when examining specific genes. Both methods demonstrated good agreement with the RNAscope score in detecting targets with medium to high expression levels, such as *PPIB* and *WFDC2*. However, for weakly expressed genes like *CCNE1*, QuantISH outperformed QuPath significantly. This variation may be attributed to differences in signal detection: signal intensity per cell area in QuantISH versus the number of estimated spots in QuPath. In particular, QuPath detects the number of transcripts in each cell but does not consider the intensity and size of each spot, which reflects the number of Z-probes bound to the target molecule. This can be problematic in degraded samples with few spots and low signal intensity. Accordingly, our results indicate that the correlation between QuantISH and QuPath improved as the expression levels increased (from score 1 to 4). Additionally, the direct comparison between QuPath-derived scores and visually estimated RNAscope scores revealed only moderate agreement, highlighting discrepancies in the detection of targets with very low (0-6) and very high (>15) numbers of spots. The difficulty for pathologists to accurately estimate the number of spots in the context of tissue heterogeneity and signal variation may contribute to this issue. Accordingly, QuPath may overestimate the number of RNA spots within formed clusters, particularly when dealing with highly expressed targets.

Another crucial point that emerged from our study is the difference between in situ hybridization and RT-ddPCR in terms of gene expression results. In agreement with (Tran et al., 2022) both automated and semi-quantitative data obtained from RNAscope showed a significant correlation with RT-ddPCR. However, RT-ddPCR was performed on bulk tissue without discrimination between stromal and cancer cells in the final quantification. Furthermore, a deeper analysis revealed that the correlation values varied depending on the level of expression and the type of gene analyzed. There was good concordance between RT-ddPCR and in situ methods for targets with medium expression levels, such as *PPIB*. Nevertheless, correlations were poorer for low and high expression levels (*CCNE1* and *WFDC2*) even after normalization. The underestimation of the number of copies in some *WFDC2* samples at extremely high expression may be due to the intrinsic dynamic range of ddPCR, which results from more than one cDNA template in each droplet. As for *CCNE1*, the poor correlation may be attributed to a combination of sample degradation, low target expression, and the Monte Carlo effect, limiting the quantification of low-amount transcripts. Sample age and the consequent nucleic acid degradation are common issues in gene expression analysis from FFPE tissues and have been reported for many methodologies, including RNAscope and RT-ddPCR. It is important to consider these factors when interpreting results (Baena-Del Valle et al., 2017; Bingham et al., 2017; Wehmas et al., 2019). In this sense, our cohort is suitable for comparing the accuracy of different methods since it includes FFPE tissue blocks with different ages and varying levels of nucleic acid degradation. As shown by our results on the housekeeping gene *PPIB*, RT-ddPCR analysis appears to be more susceptible to age block than in situ hybridization methods. This is because additional factors other than spontaneous nucleic acid degradation due to the fixation procedure can exacerbate target depletion. These factors include RNA extraction, reverse transcription efficiency, and the presence of PCR inhibitors (Ahlfen et al., 2007). Overall, these results suggest that the RNAscope assay should be preferred over RT-ddPCR for accurate gene expression analysis and that orthogonal validation of RNA biomarkers requires careful selection of samples with good nucleic acid quality, especially when dealing with low-expressed targets.

Digital image analysis can be a valid method to quantify the RNAscope signal beyond the semi-quantitative evaluation by visual assessment. Manual scoring is time-consuming and requires at least two pathologists to validate the results (Atout et al., 2022); moreover, tissue heterogeneity and variability of the RNA signal may lead to inaccurate estimates. In our study, even after accounting for samples’ degradation, both the QuantISH and QuPath approaches showed good overall agreement with the RNAscope score. In particular, QuantISH showed robust correlation results across different genes and performed well with *CCNE1* despite its relatively low and subtle gene expression. QuantISH also proved to be a more flexible and modular approach than QuPath. Although both analyses required computational resources, the modular nature and adaptability of QuantISH made it more accessible and less prone to error, unlike QuPath, which was resource-constrained and lacked flexibility, resulting in multiple crashes. It is worth noting that we analyzed the RNA hybridization signal on the QuPath software following the guidelines suggested by the RNAscope manufacturers; however, other pipelines can be used on the same program to achieve the same goal.

While our investigation has demonstrated the comparability of these approaches, it is important to note some limitations. Firstly, the quality of some spots in the TMAs was suboptimal, displaying variations in spot thickness, folded tissue, and degradation. However, both QuantISH and QuPath performed well despite these challenges. Second, our analysis did not account for the distribution of RNA expression across different areas within the spots, and we acknowledge these factors as limitations. Lastly, the selected TMA spots may not recapitulate the transcriptional heterogeneity in both cancer and stromal cells on bulk tissue samples.

In summary, our comparative assessment of various methods for quantifying gene expression in RNA-ISH TMAs has underscored the utility of fully digital techniques. Our findings demonstrate that the automated approaches can be an alternative approach for gene expression analysis, particularly highlighting QuantISH as the preferred option due to its modular and flexible software design. The potential clinical implications of adopting fully automated digital methods are promising, offering cost-effective, robust, and accessible tools. Notably, the demonstrated reliability of these automated approaches, even under suboptimal TMA conditions, suggests their prospective application in disease stratification, prognosis prediction, and tailoring personalized treatment strategies across various malignancies.

## Materials and Methods

### FFPE Samples and TMAs

FFPE tissue blocks were obtained from the National Cancer Institute of Aviano. The samples were peritoneal metastases obtained from surgical debulking of women diagnosed with stage III and IV HGSC, and already characterized in a previous study (Azzalini, Abdurakhmanova, et al., 2021; Azzalini, Barbazza, et al., 2021). Multiple 10 μm sections were cut from each tissue block for subsequent RT-ddPCR experiments. The blocks were then used to construct five tissue microarrays (TMAs) as previously described in Azzalini, Abdurakhmanova, et al., 2021 and Azzalini, Barbazza, et al., 2021. The tissue microarrays were used for the RNAscope hybridization assay.

### RNAscope assay and analysis

The in-situ detection of *PPIB*, *CCNE1*, and *WFDC2* was carried out utilizing the manual RNAscope 2.5 HD Detection Kit-Brown from Advanced Cell Diagnostics, following the manufacturer’s protocol. Briefly, 4 μm slides were cut and dried at 60°C for 1 hour. Tissue sections were subsequently deparaffinized with xylene (2×5 min) and absolute ethanol (2×1 min) at room temperature (RT), followed by a 10-minute incubation with hydrogen peroxide. Epitope retrieval was accomplished by exposing the slides to RNAscope 1x Target Retrieval Reagent for 15 min at 98°C. Tissues were then permeabilized using RNAscope protease plus (Advanced Cell Diagnostics) for 30 min at 40°C, followed by a 2-hour incubation with specific *PPIB*, *CCNE1*, and *WFDC2* RNA probes (Advanced Cell Diagnostics) at 40°C. After signal amplification, colorimetric detection was performed using DAB for 10 minutes at RT. Subsequently, slides were counterstained with hematoxylin and mounted with EcoMount solution. The total expression level, represented by the number of estimated spots per cell in tumor tissue across the entire spot, was assessed using a semi-quantitative scoring method based on the ACD scoring criteria (RNAscope score), as outlined in the RNAscope data analysis guideline (https://acdbio.com/dataanalysisguide). Representative pictures of RNAscope scoring for *CCNE1*, *PPIB*, and *WFDC2* are reported in Supplementary Figure 1.

### Imaging and image analysis

Tissue microarray slides were scanned at 40x original magnification using a 3DHISTECH Pannoramic 250 FLASH II digital slide scanner at the Genome Biology Unit supported by HiLIFE and the Faculty of Medicine, University of Helsinki, and Biocenter Finland. Subsequently, the scanned TMAs were subjected to automated RNA expression analysis utilizing QuantISH and QuPath. For the analysis, tissue spots from patients who had received neoadjuvant chemotherapy (NACT) and exhibited either no tumor cells or had missing/damaged cores have been excluded. The final dataset comprised peritoneal metastatic tissues from 55 patients diagnosed with HGSC between 2007 and 2017. The FFPE material included slices cut from five tissue microarray blocks (n = 5 TMAs), each containing an average of two cores per patient (range 1-3; each core measuring 1.2 mm), for a total of 235 tumor spots.

### QuantISH

The analysis with QuantISH adheres to the methodology outlined in the article by Jamalzadeh et al., 2022. In summary, the process involves three main steps: pre-processing, cell segmentation and classification, and RNA signal quantification.

A few modifications were implemented in the cell segmentation and classification stages to adapt the analysis to our specific dataset. A Deep Learning approach named nucleAIzer (Hollandi et al., 2020) was utilized for cell segmentation, representing a more advanced method than CellProfiler used in the original pipeline. This method was applied to each cleaned spot after pre-processing.

For cell classification, despite following the framework presented in the Jamalzadeh et al., 2022 article, we found it necessary to retrain the supervised quadratic classifier used for cell classification. This retraining involved utilizing new annotations from 580 cells, from which we extracted three features (area, mean intensity, and eccentricity) as input features for training purposes. The identified cell types include cancer, stroma, immune cells, and artifacts.

Ultimately, the RNA signal was determined as the average expression of cancer cells exclusively in each spot.

### QuPath

Various steps were executed employing QuPath software. Initially, the cores of each TMA were identified using the TMA dearrayer function. Subsequently, RNAscope images underwent analysis following the guidelines outlined in the manufacturer’s technical note “QuPath analysis guideline” (https://acdbio.com/QuPath-rna-ish-analysis). This involved adjusting stain vectors and subcellular detection parameters. To identify cells within the cores, annotations from 42 regions of interest (ROIs), each measuring 200×200 pixels, were chosen to represent TMA spots. These annotations were then employed as training images. A random tree classifier was trained to detect both cancer and non-cancer cells. The trained classifier was subsequently applied to analyze all TMA slides. Following the classification into cancer and non-cancer cells, RNA signals were identified in each core using the QuPath Subcellular Detection function. Subsequently, the quantification of the number of cancer cells, the estimated number of spots, and their mean intensity was extracted from the QuPath measurements table using Python for further comparative analyses. The QuPath script employed for the analysis is presented in the Supplementary Fig. 3.

### RT-droplet digital PCR

Total RNA was extracted from a 10-μm thick tissue section using the Maxwell® RSC RNA FFPE kit on the dedicated instrument (Promega, Madison, WI 53711-5399, USA), and 300 ng of the extracted RNA was reverse transcribed to cDNA, following previously described protocols (Bonin et al., 2019). The expression levels of *CCNE1*, *PPIB*, and *WFDC2* were subsequently determined using digital droplet PCR employing the TaqMan assay.

The correct annealing temperature for each amplicon was established through a thermal gradient. The reaction was carried out in a final volume of 20 μL, comprising 1X RT-ddPCR Supermix for Probes (no dUTP) (Bio-Rad, Hercules, CA, USA; Cat. No. 1863024), 17 ng cDNA, 18 pmol of each primer, and 5 pmol of the specific probe. A no-template sample served as a negative control in each run. The PCR reactions were partitioned into droplets using a QX200 droplet generator and transferred to a 96-well PCR plate. Endpoint PCR was conducted in a thermal cycler (iCycler, Bio-Rad) under the following conditions: enzyme activation at 95°C for 10 min; 40 cycles of denaturation at 94°C for 30 s and annealing at 56°C for 1 min at a ramp rate of 1°C/sec; enzyme inactivation at 98°C for 10 min, followed by signal stabilization at 4°C.

The fluorescence of each droplet was quantified using a QX200 droplet reader (Bio-Rad Laboratories), and the absolute concentration of the target sequence was calculated as copies/μL. The primer sequences can be found in the Supplementary Table 1.

### Statistical analysis

Statistical analyses were performed using GraphPad Prism 8.2.1 (GraphPad Software, La Jolla, CA, USA) and R version 4.3.0. For cases with multiple spots, the average expression was calculated. The distribution of the data was evaluated using the Shapiro-Wilk test to determine whether it was parametric or non-parametric. Differences between the two groups were compared using unpaired t-tests or Mann-Whitney tests for independent samples, and Wilcoxon paired tests for matched pairs. Differences among three or more groups were analyzed using the Kruskal-Wallis test for independent samples or the Friedman test for matched samples. For each method, gene expression levels were normalized concerning the housekeeping gene *PPIB* following the procedure described in gene expression data analysis guidelines of nSolver software (https://www.nanostring.com). A P-value of less than 0.05 was considered statistically significant.

## Acknowledgments

This work was supported by the European Union’s Horizon 2020 Research and Innovation Programme under grant agreements no. 667403 (HERCULES), no. 965193 (DECIDER) and 952179 (INCISIVE), the Sigrid Jusélius Foundation and the Cancer Foundation Finland (S.H. and V.A.).

## Notes

### Competing Interest Statement

The authors have declared no competing interest.

